# Machine translation of cortical activity to text with an encoder-decoder framework

**DOI:** 10.1101/708206

**Authors:** Joseph G. Makin, David A. Moses, Edward F. Chang

**Affiliations:** Center for Integrative Neuroscience/Department of Neurological Surgery, UCSF, 675 Nelson Rising Lane, San Francisco, CA 94143, USA

**Author notes:** For correspondence (JGM);, (EFC).

## Abstract

A decade after the first successful attempt to decode speech directly from human brain signals, accuracy and speed remain far below that of natural speech or typing. Here we show how to achieve high accuracy from the electrocorticogram at natural-speech rates, even with few data (on the order of half an hour of spoken speech). Taking a cue from recent advances in machine translation and automatic speech recognition, we train a recurrent neural network to map neural signals directly to word sequences (sentences). In particular, the network first encodes a sentence-length sequence of neural activity into an abstract representation, and then decodes this representation, word by word, into an English sentence. For each participant, training data consist of several spoken repeats of a set of some 30-50 sentences, along with the corresponding neural signals at each of about 250 electrodes distributed over peri-Sylvian speech cortices. Average word error rates across a validation (held-out) sentence set are as low as 7% for some participants, as compared to the previous state of the art of greater than 60%. Finally, we show how to use transfer learning to overcome limitations on data availability: Training certain components of the network under multiple participants’ data, while keeping other components (e.g., the first hidden layer) “proprietary,” can improve decoding performance—despite very different electrode coverage across participants.

## Introduction

In the last decade, brain-machine interfaces (BMIs) have transitioned from animal models into human subjects, demonstrating that some amount of motor function can be restored to tetraplegics— typically, continuous movements with two degrees of freedom (***Nuyujukian et al., 2018; Gilja et al., 2015; Jarosiewicz et al., 2015***). Although this type of control can be used in conjunction with a virtual keyboard to produce text, even under ideal cursor control (not currently achievable), the word rate would still be limited to that of typing with a single finger. The alternative is direct decoding of spoken (or attempted) speech, but heretofore such BMIs have been limited either to isolated phonemes or monosyllables (***Brumberg et al., 2009, 2011; Pei et al., 2011; Mugler et al., 2018; Stavisky et al., 2018***) or, in the case of continuous speech on moderately-sized vocabularies (about 100 words) (***Herff et al., 2015***), to decoding correctly less than 40% of words.

To achieve higher accuracies, we exploit the conceptual similarity of the task of decoding speech from neural activity to the task of machine translation, i.e., the algorithmic translation of text from one language to another. Conceptually, the goal in both cases is to build a map between two different representations of the same underlying unit of analysis. More concretely, in both cases the aim is to transform one sequence of arbitrary length into another sequence of arbitrary length— arbitrary because the lengths of the input and output sequences vary and are not deterministically related to each other. In this study, we attempt to decode a single sentence at a time, as in most modern machine-translation algorithms, so in fact both tasks map to the same kind of output, a sequence of words corresponding to one sentence. The inputs of the two tasks, on the other hand, are very different: neural signals and text. But modern architectures for machine translation learn their features directly from the data with artificial neural networks (***Sutskever et al., 2014; Cho et al., 2014b***), suggesting that end-to-end learning algorithms for machine translation can be applied with little alteration to speech decoding.

To test this hypothesis, we train one such “sequence-to-sequence” architecture on neural signals obtained from the electrocorticogram (ECoG) during speech production, and the transcriptions of the corresponding spoken sentences. The most important remaining difference between this task and machine translation is that, whereas datasets for the latter can contain upwards of a million sentences (***Germann, 2001***), a single participant in the acute ECoG studies that form the basis of this investigation typically provide no more than a few thousand. To exploit the benefits of end-to-end learning in the context of such comparatively exiguous training data, we use a restricted “language” consisting of just 30-50 unique sentences; and, in some cases, transfer learning from other participants and other speaking tasks.

## Results

Participants in the study read aloud sentences from one of two data sets: a set of picture descriptions (30 sentences, about 125 unique words), typically administered in a single session; or MOCHA-TIMIT (***Wrench, 2019***) (460 sentences, about 1800 unique words), administered in blocks of 50 (or 60 for the final block), which we refer to as MOCHA-1, MOCHA-2, etc. Blocks were repeated as time allowed. For testing, we considered only those sets of sentences which were repeated at least three times (i.e., providing one set for testing and at least two for training), which in practice restricted the MOCHA-TIMIT set to MOCHA-1 (50 sentences, about 250 unique words). We consider here the four participants with at least this minimum of data coverage.

### The Decoding Pipeline

We begin with a brief description of the decoding pipeline, illustrated in Fig. 1, and described in detail in the **Methods**. We recorded neural activity with high-density (4-mm pitch) ECoG grids from the peri-Sylvian cortices of participants, who were undergoing clinical monitoring for seizures, while they read sentences aloud. The envelope of the high-frequency component (70–150 Hz, “high-*γ*”) of the ECoG signal at each electrode was extracted with the Hilbert transform at about 200 Hz (***Dichter et al., 2018***), and the resulting sequences—each corresponding to a single sentence—passed as input data to an “encoder-decoder”-style artificial neural network (***Sutskever et al., 2014***). The network processes the sequences in three stages:

**Figure 1.**
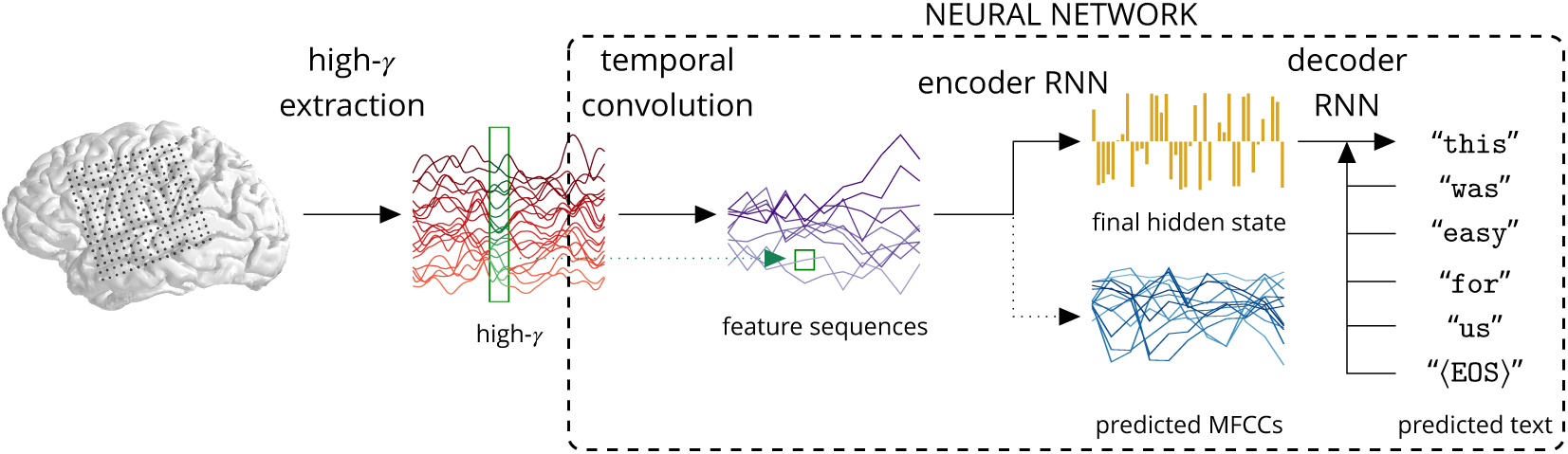
The decoding pipeline. Each participant read sentences from one of two data sets (MOCHA-TIMIT, picture descriptions) while neural signals were recorded with an ECoG array (120–250 electrodes) covering peri-Sylvian cortices. The analytic amplitudes of the high-*γ* signals (70–150 Hz) were extracted at about 200 Hz, clipped to the length of the spoken sentences, and supplied as input to an artificial neural network. The early stages of the network learn temporal convolutional filters that, additionally, effectively downsample these signals. Each filter maps data from twelve-sample-wide windows across all electrodes (e.g., the green window shown on the example high-*γ* signals in red) to single samples of a feature sequence (highlighted in the green square on the blue feature sequences); then slides by twelve input samples to produce the next sample of the feature sequence; and so on. One hundred feature sequences are produced in this way, and then passed to the *encoder* RNN, which learns to summarize them in a single hidden state. The encoder RNN is also trained to predict the MFCCs of the speech audio signal that temporally coincide with the ECoG data, although these are not used during testing (see text for details). The final encoder hidden state initializes the *decoder* RNN, which learns to predict the next word in the sequence, given the previous word and its own current state. During testing, the previous *predicted* word is used instead.

1. *Temporal convolution*: A fully connected, feed-forward network cannot exploit the fact that similar features are likely to recur at different points in the sequence of ECoG data. For example, the production of a particular phoneme is likely to have a similar signature at a particular electrode independently of when it was produced (***Bouchard et al., 2013***). In order to learn efficiently such regularities, the network applies the same temporally brief filter (depicted with a rectangle across high-*γ* waveforms in Fig. **1**) at regular intervals (“strides”) along the entire input sequence. Setting the stride greater than one sample effectively downsamples the resulting output sequence. Our network learns a set of such filters, yielding a set of filtered sequences effectively downsampled to 16 Hz.
2. *Encoder recurrent neural network* (RNN): The downsampled sequences are consumed seriatim by an RNN. That is, at each time step, the input to the encoder RNN consists of the current sample of each of the downsampled sequences, as well as its own previous state. The *final* hidden state (yellow bars in Fig. 1) then provides a single, high-dimensional encoding of the entire sequence, independent of its length. In order to guide the encoder toward useful solutions during training, we also require it to predict, at each time step, a representation of the speech audio signal, the sequence of Mel-frequency cepstral coefficients (MFCCs; see **Methods**).
3. *Decoder RNN*: Finally, the high-dimensional state must be transformed back into another sequence, this time of words. A second RNN is therefore initialized at this state, and then trained to emit at each time step either a word or the end-of-sequence token—at which point decoding is terminated. At each step in the output sequence, the decoder takes as input, in addition to its own previous hidden state, either the preceding word in the actual sentence uttered by the participant (during the model-training stage), or its own predicted word at the preceding step (during the testing stage). The use of words for targets stands in contrast to previous attempts at speech decoding, which target phonemes (***Brumberg et al., 2009, 2011; Pei et al., 2011; Herff et al., 2015; Mugler et al., 2018; Stavisky et al., 2018***).

The entire network is simultaneously trained to make the encoder emit values close to the target MFCCs, and the decoder assign high probability to each target word. Note that the MFCC-targeting provides an “auxiliary loss,” a form of multi-task learning (***Caruana, 1997; Szegedy et al., 2015***): Its purpose is merely to guide the network toward good solutions to the word-sequence decoding problem; during testing, the MFCC predictions are simply discarded, and decoding is based entirely on the decoder RNN’s output. All training proceeds by stochastic gradient descent via backpropagation (***Rumelhart et al., 1986***), with dropout (***Srivastava et al., 2014***) applied to all layers. (A more mathematical treatment of the decoder appears in the **Methods**.)

### Decoding Performance

Here and throughout, we quantify peformance with the average (across all tested sentences) word error rate (WER), i.e., the minimum number of deletions, insertions, and substitutions required to transform the predicted sentence into the true sentence, normalized by the length of the latter. Thus the WER for perfect decoding is 0%, and for erroneous decoding is technically unbounded, although in fact can be capped at 100% simply by predicting empty sentences. For reference, in speech transcription, word error rates of 5% are professional-level (***Xiong et al., 2017***), and 20-25% is the outer bound of acceptable performance (***Munteanu et al., 2006***). It is also the level at which voice-recognition technology was widely adopted, albeit on much larger vocabularies (***Schalkwyk et al., 2010***)

We begin by considering the performance of the encoder-decoder framework for one example participant speaking the 50 sentences of MOCHA-1 (50 sentences, about 250 unique words), Fig. 2A. The WER for the participant shown in Fig. 2A is approximately 7%. The previous state-of-the-art WER for speech decoding is 60%, and operated with a smaller (100-word) vocabulary (***Herff et al., 2015***).

**Figure 2.**
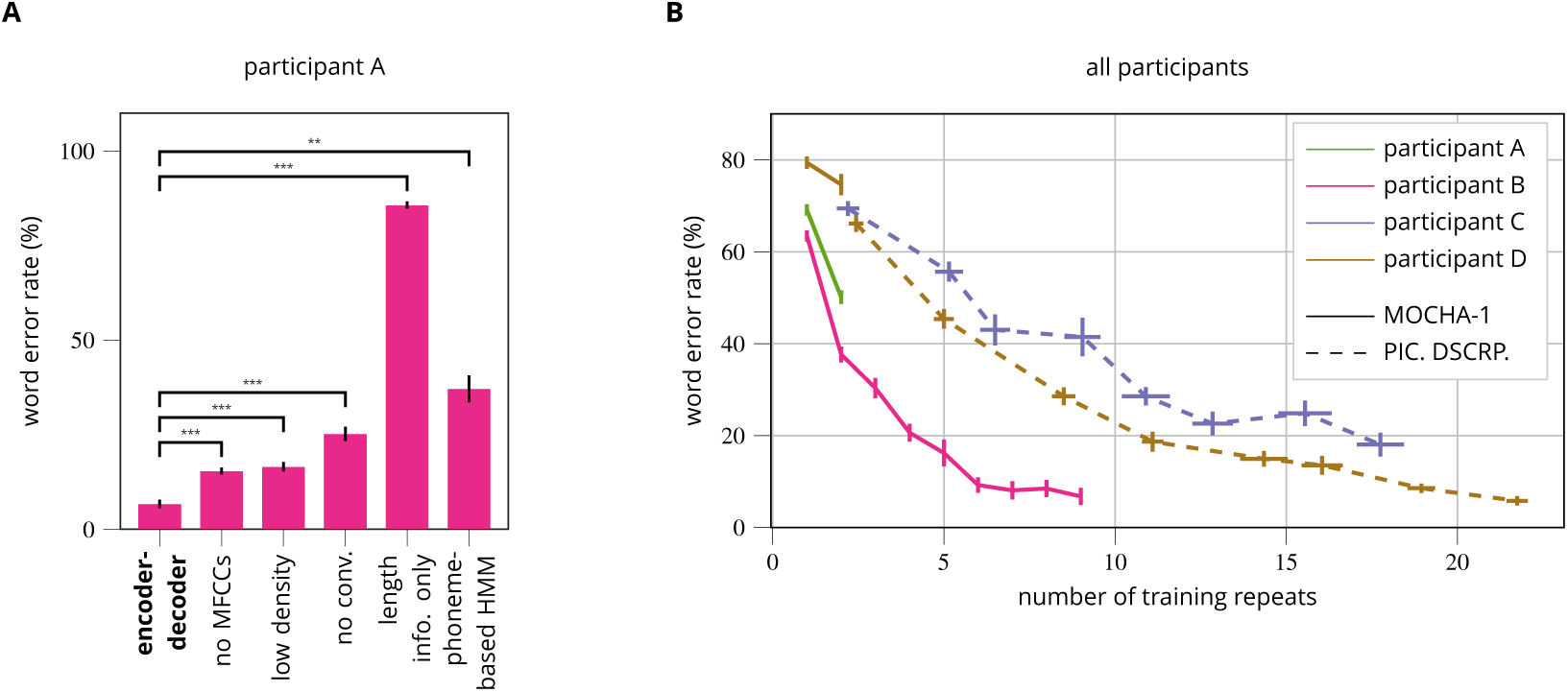
Word error rates (WERs) of the decoded sentences. **(A)** WERs for one participant under the encoder-decoder (first bar), four crippled variants thereof (bars 2-5), and a state-of-the-art phoneme-based Viterbi decoder (“phoneme-based HMM”). Abbreviations: “no MFCCs”: trained without requiring the encoder to predict MFCCs; “low density”: trained and tested on simulated lower-density grid (8-mm rather than 4-mm spacing); “no conv.”: the network’s temporal convolution layer is replaced with a fully connected layer; “length info. only”: the input ECoG sequences are replaced with Gaussian noise—but of the correct length. Whiskers indicate standard errors of the mean WERs across 30 networks trained from scratch and evaluated on randomly selected held-out blocks (except the Viterbi decoder which did not vary across retrainings and was therefore simply leave-one-out cross-validated under the ten available blocks). Significance is indicated by stars (∗= *p*< 0.05, ∗∗= *p*< 0.005, ∗∗∗= *p*< 0.0005). **(B)** For four different participants, WER as a function of the number of repeats of the sentence sets used for training, i.e. the number of training tokens for each sentence type. Results for MOCHA-1 (50 sentence types; see text for detail) are shown in solid lines (pink, green, brown); for the picture descriptions (“PIC. DSCRP.”; 30 sentence types), in dashed lines (blue, brown). Note that participant D (brown) read from both sets. The endpoint of the pink curve corresponds to the first bar of **(A)**. Whiskers indicate standard errors of the mean WERs (vertical) and mean number of repeats (horizontal) across ten networks trained from scratch and evaluated on randomly selected held-out blocks. (The number of repeats varies because data were divided on the basis of blocks, which vary slightly in length.)

Since all training and testing sentences belong to the same set of 50, it is also possible to compare performance against a *sentence classifier* (as opposed to word-by-word decoder). Here we compare against a state-of-the-art phoneme-based classifier for speech production (***Moses et al., in press***). Briefly, each of the 50 MOCHA-1 sentences was assigned its own hidden Markov model (HMM), whose emissions are neural activity and whose hidden state can transition only through the phonemes of that sentence (including self transitions). Test sequences of ECoG data were classified by choosing the HMM—and corresponding MOCHA-1 sentence—whose most-likely phoneme sequence (Viterbi path) had the highest probability across all 50 HMMs. See the **Methods** for details. The WER (Fig. 2A, “phoneme-based HMM”) for this model, approximately 37%, is more than five times that of the encoder-decoder network.

What accounts for the superior performance of the encoder-decoder network? To quantify the contributions of its various elements, we systematically remove or cripple them, and retrain networks from scratch. The second bar shows performance when MFCCs are not targeted during training. Thus, where speech audio is not available, as may well be the case for a candidate for a speech prosthesis, error rate is more than 100% greater—although again within the usable range. The third bar shows performance on data that have been spatially downsampled in order to simulate lower-density ECoG grids. Specifically, we simply discarded every other channel along both dimensions of the grid, leaving just one quarter of the channels, i.e. nominally 64 instead of 256. Performance is similar to the model trained without speech-audio targeting, but notably superior to previous attempts at speech decoding and still within the usable range, showing the importance of the algorithm in addition to high-density grids. Next, we consider a network whose input layer is fully connected, rather than convolutional (fourth bar). Word error rates quadruple. Note that the temporal convolution in our model also effectively downsamples the signal by a factor of twelve (see **The Decoding Pipeline** above), bringing the length of the average sequence seen by the encoder RNN down from about 450 to about 40 samples. And indeed, our exploratory analyses showed that some of the performance lost by using fully connected input layers can be recovered simply by downsampling the high-*γ* activity before passing it to them. Thus the decrease in performance due to removing temporal convolution may be explained in part by the difficulty encoder-decoder networks have with long input sequences (***Cho et al., 2014a***).

Recall that the endpoints of each ECoG sequence fed to the encoder-decoder were determined by the endpoints of the corresponding speech audio signal. Thus it might seem possible for the network to have learned merely the (approximate) length of each unique sentence in MOCHA-1, and then during testing to be simply classifying them on this basis, the decoder RNN having learned to reconstruct individual sentences from an implicit class label. To show that this is not the case, we replace each sample of ECoG data with (Gaussian) noise, retrain encoder-decoder networks, and re-test. Performance is much worse than that of any of the decoders (WERs of about 86%; “length info. only” bar in Fig. 2A).

Next we consider how many data are required to achieve high performance. Fig. 2B shows WER for all four participants as a function of the number of repeats of the training set (solid lines: MOCHA-1/50 sentences; dashed lines: picture descriptions/30 sentences) used as training data for the neural networks. We note that for no participant did the total amount of training data exceed 40 minutes in total length. When at least 15 repeats were available for training, WERs could be driven below 20%, the outer bound of acceptable speech transcription, although in the best case (participant B/pink) only 5 repeats were required. On two participants (participant B/pink, participant D/brown), training on the full training set yielded WERs of about 7%, which is approximately the performance of professional transcribers for spoken speech (***Xiong et al., 2017***).

### Transfer Learning

In Fig. 2B we included two participants with few training repeats of the MOCHA sentences (participant A/green, participant D/brown) and, consequently, poor decoding performance. Here we explore how performance for these participants can be improved with *transfer learning* (***Pratt et al., 1991; Caruana, 1997***), that is, by training the network on a related task, either in parallel with or before training on the decoding task at hand, namely the MOCHA-1 sentence set.

We begin with participant A, who spoke only about four minutes of the MOCHA-1 data set (i.e., two passes through all 50 sentences, not counting the held-out block on which performance was evaluated). The first bar of Fig. 3A (“encoder-decoder”) shows WER for encoder-decoder networks trained on the two available blocks of MOCHA-1 (corresponding to the final point in the green line in Fig. 2B), which is about 48%. Next we consider performance when networks are first *pre-trained* (see **Methods** for details) on the more plentiful data for participant B (ten repeats of MOCHA-1). Indeed, this transfer-learning procedure decreases WER by about 15% (from the first to the second bar, “subject TL,” of Fig. 3A; the improvement is significant under a Wilcox signed-rank test, Holm-Bonferroni corrected for multiple comparisons, with *p*< 0.001).

**Figure 3.**
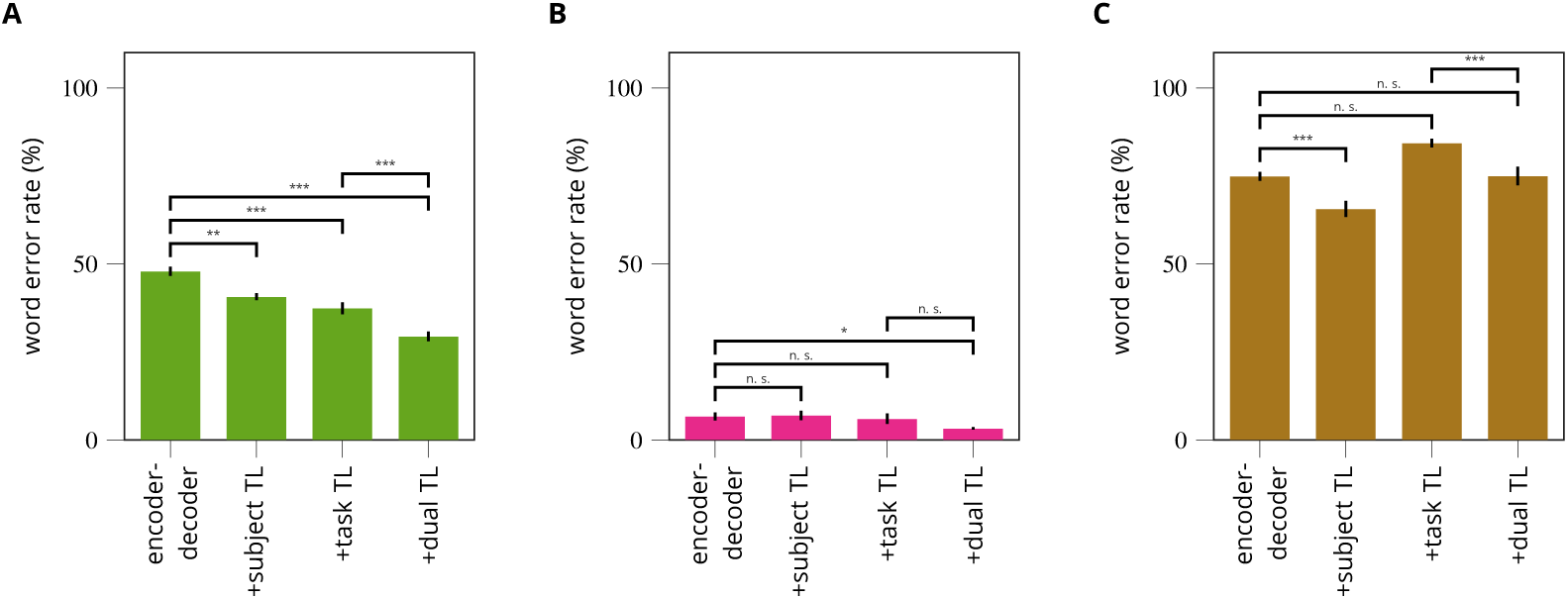
Word error rates (WERs) of the decoded MOCHA-1 sentences for encoder-decoder models trainer with *transfer learning*. Each subfigure corresponds to a subject (color code as in Fig. 2). The four bars in each subfigure show WER without transfer learning (“encoder-decoder,” as in the final points in Fig. 2B), with cross-subject transfer learning (“+subject TL”), with training on sentences outside the test set (“+task TL”), and with both forms of transfer learning (“+dual TL”). **(A)** Participant A, with pretraining on participant B/pink (second and fourth bars). **(B)** Participant B, with pretraining on participant A/green (second and fourth bars). **(C)** Participant D, with pretraining on participant B/pink (second and fourth bars). Significance is indicated by stars (∗= *p*< 0.05, ∗∗= *p*< 0.005, ∗∗∗= *p*< 0.0005, n.s.= not significant).

We have so far excluded blocks of MOCHA-TIMIT beyond the first 50 (MOCHA-1), since we were unable to collect a sufficient number of repeats for training and testing. However, the data from these excluded blocks may nevertheless provide sub-sentence information that is useful for decoding MOCHA-1, in particular word-level labels as well perhaps as information about lower-level features in the ECoG data. To test this hypothesis, we extend the training set to include also the rest of the MOCHA sentences spoken by participant A—namely, two repeats of MOCHA-2 through MOCHA-9, which together comprise 410 unique sentences (disjoint from MOCHA-1); train from scratch on this complete set of MOCHA-TIMIT; and test again on MOCHA-1. This cross-task training decreases WER by 22% over the baseline (from the first to the third bar, “task TL,” of Fig. 3A; *p*≪ 0.001). This result is particularly important because it shows that the encoder-decoder is not merely classifying sentences (in the encoder) and then reconstructing them (in the decoder), without learning their constituent parts (words), in which case the decoding scheme would not generalize well. Instead, the network is evidently learning sub-sentence information.

Finally, we consider a combined form of transfer learning, in which encoder-decoder networks are pre-trained on all MOCHA-TIMIT data for participant B (an additional single set of MOCHA-2 through MOCHA-9); then trained on all MOCHA-TIMIT data for participant A; and then tested as usual on a held-out block of MOCHA-1 for participant A. This “dual transfer learning” decreases WER an additional 21% (Fig. 3A, third and fourth bars, *p*≪ 0.001), or a full 39% over baseline performance.

Although participant B has less room for improvement, we consider whether it is nevertheless possible to decrease WER with transfer learning (Fig. 3B). Cross-subject (from participant A) transfer learning alone (“subject TL”) does not improve performance, probably because of how few blocks it adds to the training set (just three, as opposed to the ten that are added by transfer in the reverse direction). The improvement under cross-task transfer learning (“task TL”) is not significant, again presumably because it increases the number of training blocks only by a factor of two, as opposed to the factor of almost ten for participant A (Fig. 3A). Used together, however, cross-task and cross-subject transfer reduce WER by 46% over baseline (first and fourth bars; *p* = 0.03 after correcting for multiple comparisons)—indeed, down to about 3%.

For the participant with the worst performance on the MOCHA-TIMIT data, participant D (see again Fig. 3C, solid brown line), adding the rest of the MOCHA sentences to the training set does not improve results, perhaps unsurprisingly (Fig. 3C, “task TL”). However, cross-subject transfer learning (from participant B into participant D) again significantly improves decoding (*p* < 0.001). Finally, for the two participants reading picture descriptions, subject transfer learning does not improve results.

### Anatomical Contributions

To determine what areas of the cortex contribute to decoding in the trained models, we compute the derivative of the loss function with respect to the electrode activities across time. These values measure how the loss function would be changed by small changes to the electrode activities, and therefore the relative importance of each electrode. (A similar technique is used to visualize the regions of images contributing to object identification by convolutional neural networks (***Simonyan et al., 2013***).) Under the assumption that positive and negative contributions to the gradient are equally meaningful, we compute their norm (rather than average) across time and examples, yielding a single (positive) number for each electrode. See the **Methods** for more details.

Fig. 4 shows, for each of the four participants, the distribution of these contributions to decoding within each anatomical area. (For projections onto the cortical surface, see Fig. S1 in the Supplementary Material.) In all subjects with left-hemisphere coverage (B/pink, C/blue, D/brown), strong contributions are made by ventral sensorimotor cortex (vSMC), as expected from the cortical area most strongly associated with speech production (***Conant et al., 2018***). However, there is heterogeneity across participants, some of which can perhaps be explained by differing grid coverage. For participant D, who had reduced motor coverage due to malfunctioning electrodes (along the missing strips seen in Fig. S1), decoding relies heavily on superior temporal areas, presumably decoding auditory feedback, either actual or anticipated. In the single patient with right-hemisphere coverage (A/green), the inferior frontal gyrus, in particular the pars triangularis, contributed disproportionately to decoding. (This participant, like the others, was determined to be left-hemisphere language-dominant, so these highly contributing electrodes cover the *non-dominant* homologue of Broca’s area.) Somewhat surprisingly, in the patient for whom the best decoding results were obtained, B/pink, the supramarginal gyrus of posterior parietal cortex makes a large and consistent contribution.

**Figure 4.**
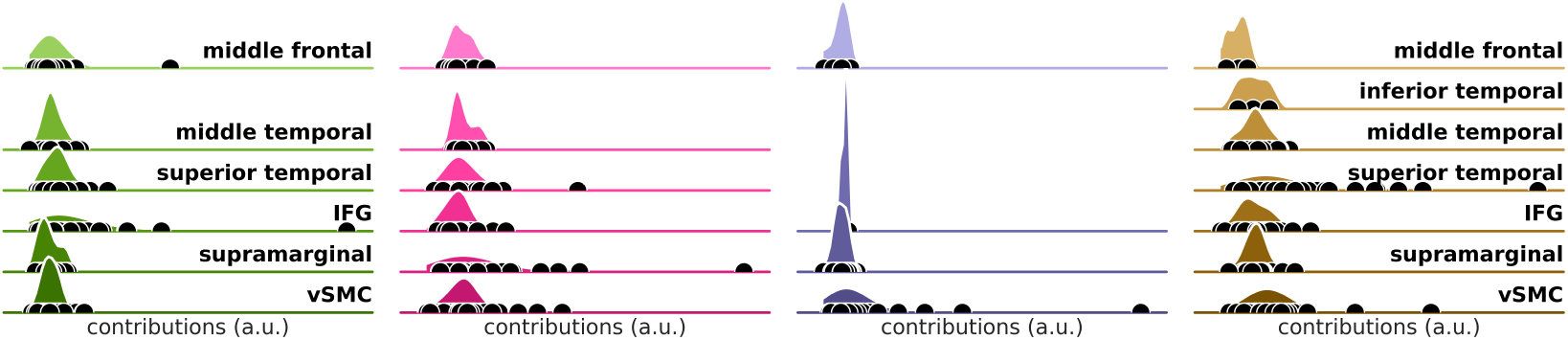
The contributions of each anatomical area to decoding, as measured by the gradient of the loss function with respect to the input data (see main text for details). The contributions are broken down by participant, with the same color scheme as throughout (cf. Fig. 2). Each shaded area represents a kernel density estimate of the distribution of contributions of electrodes in a particular anatomical area; black dots indicate the raw contributions. The scale and “zero” of these contributions were assumed to be incomparable across participants and therefore all data were rescaled to the same interval for each subject (smallest contribution at left, largest contribution at right). Missing densities (e.g., temporal areas in the participant C/blue) correspond to areas with no grid coverage. IFG: inferior frontal gyrus; vSMC: ventral sensorimotor cortex.

## Discussion

We have shown that spoken speech can be decoded reliably from ECoG data, with WERs as low as 3% on data sets with 250-word vocabularies. But there are several provisos. First, the speech to be decoded was limited to 30-50 sentences. The decoder learns the structure of the sentences and uses it to improve its predictions. This can be seen in the errors the decoder makes, which frequently include pieces or even the entirety of other valid sentences from the training set (see Table 1). Although we should like the decoder to learn and to exploit the regularities of the language, it remains to show how many data would be required to expand from our tiny languages to a more general form of English.

**Table 1.**
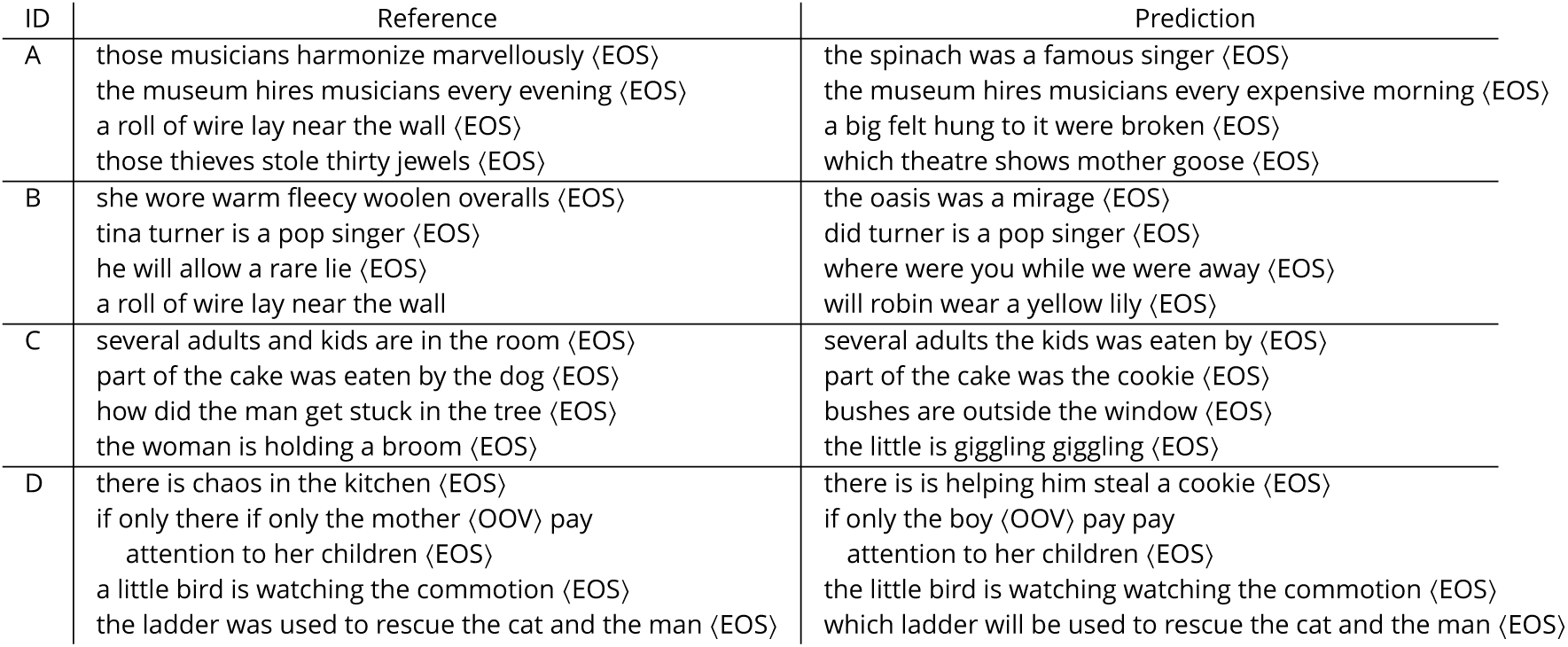
Example incorrectly decoded sentences (“Prediction,” right) and the actual sentence spoken (“Reference,” left) for participants A–D. ID = participant ID, ⟨EOS⟩ = the end-of-sequence token, ⟨OOV⟩ = out-of-vocabulary token

On the other hand, the network is not merely classifying sentences, since performance is improved by augmenting the training set even with sentences *not* contained in the testing set (Fig. 3A). This result is critical: it implies that the network has learned to identify words, not just sentences, from ECoG data, and therefore that generalization to decoding of *novel* sentences is possible. Indeed, where data are plentiful, encoder-decoder models have been shown to learn very general models of English (***Bahdanau et al., 2014***). And, as we have seen, the number of data required can be reduced by pre-training the network on other participants—even when their ECoG arrays are implanted in different hemispheres (Fig. 3, Fig. S1). In principle, transfer learning could also be used to acquire a general language model without any neural data at all, by pre-training an encoder-decoder network on a task of translation to, or autoencoding of, the target language (e.g., English)—and then discarding the encoder.

We attribute the success of this decoder to three major factors. First, recurrent neural networks with long short-term memory are known to provide state-of-the-art information extraction from complex sequences, and the encoder-decoder framework in particular has been shown to work well for machine translation, a task analogous to speech decoding. Furthermore, the network is trained end-to-end, obviating the need to hand-engineer speech-related neural features about which our knowledge is quite limited. This allows the decoder to be agnostic even about which cortical regions might contribute to speech decoding.

Second, the most basic labeled element in our approach is the word, rather than the phoneme as in previous approaches. Here the trade-off is between coverage and distinguishability: Far fewer phonemes than words are required to cover the space of English speech, but individual phonemes are shorter, and therefore less distinguishable from each other, than words. In fact, the production of any particular phoneme in continuous speech is strongly influenced by the phonemes preceding it (“coarticulation”), which decreases their distinguishability still further (or, equivalently, reduces coverage by requiring parsing in terms of biphones, triphones, or even quinphones). At the other extreme, English sentences are even more distinguishable than words, but their coverage is much worse. Of course, in this study we have limited the language to just a few hundred words, artificially reducing the cost of poor coverage. But our results suggest that expanding the amount of data beyond 30 minutes will allow for an expansion in vocabulary and flexibility of sentence structure. We also note that even a few hundred words would be quite useful to a patient otherwise unable to speak at all. Finally, the use of words rather than phonemes may also make possible access to semantic and lexical representations in the cortex.

Third and finally, decoding was improved by modifying the basic encoder-decoder architecture (***Sutskever et al., 2014***) in two ways: adding an auxiliary penalty to the encoder that obliges the middle layer of the RNN to predict the MFCCs of the speech audio; and replacing the fully-connected feedforward layers with temporal-convolution layers, which also effectively downsamples the incoming ECoG signals by a factor of about ten. In fact, very recent work in machine learning has shown that RNNs can sometimes be replaced entirely with temporal-convolution networks, with superior results (***Bai et al., 2018***)—a promising avenue for future improvements to the decoder presented here.

We have emphasized the practical virtue of neural networks learning their own features from the data, but it comes at a scientific price: the learned features—in this case, neural activity in different brain regions—can be difficult to characterize. This is an open research area in machine learning. What we can say (cf. Fig. 4) is that all anatomical areas covered by the ECoG grid appear to contribute to decoding, with a generally large contribution from the ventral sensorimotor cortex (vSMC) (***Conant et al., 2018***). Nevertheless, there are important variations by subject, with right pars triangularis, the supramarginal gyrus, and the superior temporal gyrus contributing strongly in, respectively, participant A/green, participant B/pink, and participant D/brown. We also note that the contributions of vSMC are disproportionately from putatively sensor, i.e. postcentral, areas (Fig. S1). It is possible that this reflects our choice to clip the high-*γ* data used for training at precisely the sentence boundaries, rather than padding the onsets: Truncating the speech-onset commands may have reduced the salience of motor features relative to their sensory consequences (including efference copy and predicted tactile and proprioceptive responses (***Tian and Poeppel, 2010***)).

To investigate the *kinds* of features being used, one can examine the patterns of errors produced. However, these are not always indicative of the feature space used by the network, whose errors often involve substitution of phrases or even whole sentences from other sentences of the training set (a strong bias that presumably improves decoding performance overall by guiding decoded output toward “legitimate” sentences of the limited language). Nevertheless, some examples are suggestive. There appear to be phonemic errors (e.g., in Table 1, “robin wear” for “roll of wire,” “theatre” for “thieves,” “did” for “tina”), as expected, but also semantic errors—for example, the remarkable series of errors for “those musicians harmonize marvellously,” by different models trained on the data from participant A, in terms of various semantically related but lexically distinct sentences (“the spinach was a famous singer,” “tina turner those musicians harmonize singer,” “does turner ⟨OOV⟩ increases”). Since the focus of the present work was decoding quality, we do not pursue questions of neural features any further here. But these examples nevertheless illustrate the utility of powerful decoders in revealing such features, and we consider a more thorough investigation to be the most pressing future work.

Finally, we consider the use of the encoder-decoder framework in the context of a brain-machine interface, in particular as a speech prosthesis. The decoding stage of the network already works in close to real time. Furthermore, in a chronically implanted participant, the amount of available training data will be orders of magnitude greater than the half hour or so of speech used in this study, which suggests that the vocabulary and flexibility of the language might be greatly expandable. On the other hand, MFCCs may not be available—the participant may have already lost the ability to speak. This will degrade performance, but not insuperably (Fig. 2A). Indeed, without MFCCs, the only data required beyond the electrocorticogram and the text of the target sentences is their start and end times—a distinct advantage over decoders that rely on phoneme transcription. A more difficult issue is likely to be the changes in cortical representation induced by the impairment or by post-impairment plasticity. Here again the fact that the algorithm learns its own features—and indeed, learns to use brain areas beyond primary motor and superior temporal areas—make it a promising candidate.

## Methods

The participants in this study were undergoing treatment for epilepsy at the UCSF Medical Center. Electrocorticograpic (ECoG) arrays were surgically implanted on each patient’s cortical surface in order to localize the foci of their seizures. Prior to surgery, the patients gave written informed consent to participate in this study, which was executed according to protocol approved by the UCSF Committee on Human Research. All four participants of this study were female, right-handed, and determined to be left-hemisphere language-dominant.

### Task

Participants read sentences aloud, one at a time. Each sentence was presented briefly on a computer screen for recital, followed by a few seconds of rest (blank display). Two participants (A and B) read from the 460-sentence set known as MOCHA-TIMIT (***Wrench, 2019***). These sentences were designed to cover essentially all the forms of coarticulation (connected speech processes) that occur in English, but are otherwise unremarkable specimens of the language, averaging 9±2.3 words in length, yielding a total vocabulary of about 1800 unique words. Sentences were presented in blocks of 50 (or 60 for the ninth set), within which the order of presentation was random (without replacement). The other two participants (C and D) read from a set of 30 sentences describing three (unseen) cartoon drawings, running 6.4±2.3 words on average, and yielding a total vocabulary of about 125 words; see Table 2. A typical block of these “picture descriptions” consisted of either all 30 sentences or a subset of just 10 (describing one picture).

**Table 2.** The picture descriptions read by participants C and D. N.B. that patients did not view the pictures.

| pic. # | sentence |
| --- | --- |
| 1 | part of the cake was eaten by the dog<br>several adults and kids are in the room<br>the little boy is crying because the dog ate his cake<br>the mother is angry at her pet dog<br>under the sofa is a hiding dog<br>the woman is holding a broom<br>there is a partially eaten cake on the large table<br>four candles are lit on the cake<br>the guests arrived with presents<br>the child is turning four years old |
| 2 | while falling the boy grabs a cookie<br>the boy is reaching for the cookie jar<br>there is chaos in the kitchen<br>water is overflowing from the sink<br>if only the mother could pay attention to her children<br>the stool is tipping over<br>the little girl is giggling<br>his sister is helping him steal a cookie<br>bushes are outside the window<br>i think their water bill will be high |
| 3 | the firemen are coming to the rescue<br>the girl was riding a tricycle<br>which ladder will be used to rescue the cat and the man<br>the cat does not want to come off the tree branch<br>in the tree there is a cat, a man, and a bird<br>a dog is barking at the man in the tree<br>a little bird is watching the commotion<br>worried by the dog the man considers jumping<br>the cat doesnt seem interested in coming down<br>how did the men get stuck in the tree |

The reading of these blocks was distributed across several days. The number of passes through the entire set depended on available time and varied by patient. The breakdown is summarized in Table 3.

**Table 3.** Data sets for training and testing, broken down by participant. MT-1 = MOCHA-TIMIT, first set of 50 sentences; MT-* = MOCHA-TIMIT, full set of 460 sentences for training, first set of 50 for testing; PD = picture descriptions. The numbers of tokens are given for a (typical) fold of cross validation but in practice could vary slightly because the cross-validation procedure partitioned the data by blocks rather than sentences. The numbers of sentence types are nominal, i.e. were not increased to reflect (rare) participant misreadings.

| participant |  |  | A |  | B |  | C | D |  |  |
| --- | --- | --- | --- | --- | --- | --- | --- | --- | --- | --- |
| data set |  |  | MT-1 | MT-* | MT-1 | MT-* | PD | MT-1 | MT-* | PD |
| training | sentence | types | 50 | 460 | 50 | 460 | 30 | 50 | 460 | 30 |
|  |  | tokens | 100 | 924 | 450 | 860 | 559 | 100 | 909 | 740 |
|  | word | types | 239 | 1787 | 240 | 1787 | 122 | 238 | 1745 | 123 |
|  |  | tokens | 610 | 6897 | 2740 | 5890 | 5453 | 607 | 6729 | 7292 |
| validation | sentence | types | 50 | 50 | 50 | 50 | 30 | 50 | 50 | 30 |
|  |  | tokens | 50 | 50 | 50 | 50 | 60 | 50 | 50 | 82 |
|  | word | types | 238 | 238 | 239 | 239 | 122 | 230 | 230 | 122 |
|  |  | tokens | 304 | 304 | 303 | 303 | 592 | 302 | 302 | 809 |

### Data collection and pre-processing

#### Neural Data

The recording and pre-processing procedures have been described in detail elsewhere (***Dichter et al., 2018***), but we repeat them briefly here. Participants were implanted with 4-mm-pitch ECoG arrays in locations chosen for clinical purposes. Three participants (A, B, D) were implanted with 256-channel grids over peri-Sylvian cortices; the remaining participant was implanted with a 128-channel grid located dorsal to the Sylvian fissure, primarily over premotor, motor, and primary sensory cortices (see Fig. S1). Grids were implanted over the left hemisphere of all patients except participant A. Analog ECoG signals were amplified and then digitized at about 3 kHz. These digital signals were then anti-aliased (low-pass filtered at 1.5 kHz), notch filtered to remove line noise, and then anti-aliased and downsampled to about 400 Hz. Next, the mean (across channels) signal was removed from all channels (“common-average referencing”). Then the analytic amplitudes (at each channel) were extracted in each of eight adjacent frequency bands between 70 and 150 Hz, averaged across bands, and downsampled to about 200 Hz. (Two participants’ data were downsampled by an integer divisor to 190 Hz, whereas the other two sets were re-sampled by a non-integer divisor to precisely 200 Hz.) Finally, these analytic amplitudes were z-scored on the basis of a 30-second sliding window, yielding the “high-*γ*” signals dicussed in the main text.

#### Speech Transcriptions

Speech was transcribed at the word level by hand or, where aided by speech-to-text software, with manual correction. Participants did not always read the sentences correctly, so the actual spoken vocabulary was generally a superset of the nominal vocabularies of the MOCHA-TIMIT or picture-description sentence sets, including non-words (false starts, filled pauses, mispronunciations, and the like). Nevertheless, the decoder represents words with a “one-hot” encoding, and consequently requires a fixed-size vocabulary. In order to allow the decoder to generalize to new participants engaged in the same task (viz., producing sentences from either MOCHA-TIMIT or the picture descriptions), all words not in the nominal sets (less than 1% of the total) were replaced with a single out-of-vocabulary token prior to their use as training data.

Sentence onset and offset times were manually extracted and used to clip the neural data into sentence-length sequences.

#### Speech Audio Signal

The speech audio of the participants was recorded simultaneously with the neural data at about 24 kHz with a dedicated microphone channel, and time aligned.

Mel-frequency cepstral coefficients (MFCCs) are features commonly extracted from speech audio for the purpose of rendering linguistic (phonemic) content more perspicuous. Briefly, the coefficients at each “frame” characterize (the logarithm of) the local power spectrum in log-spaced bins, a spacing that reflects the frequency discrimination of human hearing. Following typical practice, we used the leading 13 coefficients (of the discrete cosine transform), replacing the first of these with the log of the total frame energy. We extracted MFCCs in Python with the `python_speech_features` package (***Lyons, 2018***), using 20-ms sliding frames with a slide of 1/*F*_sampling_, where *F*_sampling_ is the sampling rate of the high-*γ* data (about 200 Hz).

### The Network

#### High-Level Description

The encoder-decoder is an artificial neural network—essentially an extremely complicated, parameterized function that is constructed by composition of simple functions, and “trained” by changing those parameters so as incrementally to decrease a penalty on its outputs. In our case, the input, outputs, and penalties for a single sentence are:

- **input**: the sequence of high-*γ* vectors (with the length of each vector the number of recording electrodes) recorded during production of the sentence;
- **outputs**: the sequence of predicted Mel-frequency cepstral coefficients (MFCCs) extracted from the speech audio signal, and the sequence of predicted words;
- **penalties**: the deviations of the predicted from the *observed* sequences of MFCCs and words.

The deviations are quantified in terms of cross entropy. For each word in the sequence, cross entropy is (proportional to) the average number of yes/no questions that would be required to “guess” correctly the true word, given the output (predicted probabilities) of the decoder. For each element (vector) of the MFCC sequence, which is assumed to be normally distributed, the cross entropy is just the mean square error between the observed and predicted vectors (plus a constant term). At each step of the training procedure, the cross entropies are computed over a randomly chosen subset of all sentences, and the parameters (weights) of the network are changed in the direction that decreases these penalties. Note that we do not actually use the predicted MFCCs during the testing phase: the point of training the network to predict the speech audio signal is simply to guide the network toward solutions to the primary goal, predicting the correct sequence of words (***Caruana, 1997***).

#### Mathematical Description

We now describe and justify this procedure more technically. Notational conventions are standard: capital letters for random variables, lowercase for their instantiations, boldface italic font for vectors, and italic for scalars. We use angle brackets, ⟨·⟩, strictly for sample averages (as opposed to expectation values). For empirical probability distributions of data generated by “the world,” we reserve *p*, and for distributions under models, *q*. The set of all parameters of the model is denoted Θ.

Consider the probabilistic graphical model in Fig. 5A. Some true but unknown relationships (denoted by the probability distribution *p*) obtain between the sequences of spoken words, 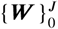, corresponding audio waveforms, 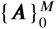, and contemporaneous neural activity, 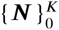 (commands to the articulators, neurolinguistic representations, efference copy, auditory feedback, etc.).^1^ The task of the network is to predict (provide probability distribution *q* over) each MFCC and word sequence, given just a neural sequence as input. The training procedure thus aims to bring *q* closer to *p*, or more precisely to minimize the conditional KL divergences,

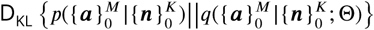

and

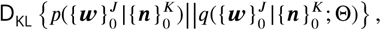

averaged under the observed data 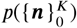, by improving the parameters Θ. This is the standard formulation for fitting probabilistic models to data.

The minimization can equivalently be written in terms of cross entropies (by dropping the entropy terms from the KL divergences, since they do not depend on the parameters), which can be further simplified by assuming specific forms for the conditional distributions over MFCCs and words. In particular, we assume that at each step *m* in the sequence, the deviations of the observed vector of MFCCs, ***a***_*m*_, from the model predictions, 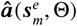, are normally distributed and conditionally independent of all other sequence steps:

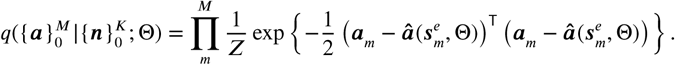

(For simplicity we let the covariance matrix be the identity matrix here, but it does not affect the minimization.) The prediction 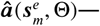 the vector output of the encoder recurrent neural network (RNN) at step *m*—depends on the entire sequence of neural data only by way of the encoder state at step *m*:

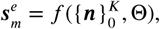

where the function *f* is given by the encoder RNN. (In the networks discussed in the main text, *f* is a three-layer bidirectional network of LSTM cells; see the discussion of the architecture below.) The cross entropy for the MFCC sequences then becomes

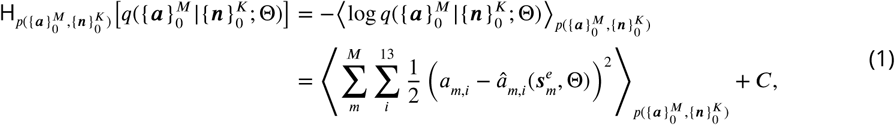

where the inner sum is over all 13 coefficients used at each step.

**Figure 5.**
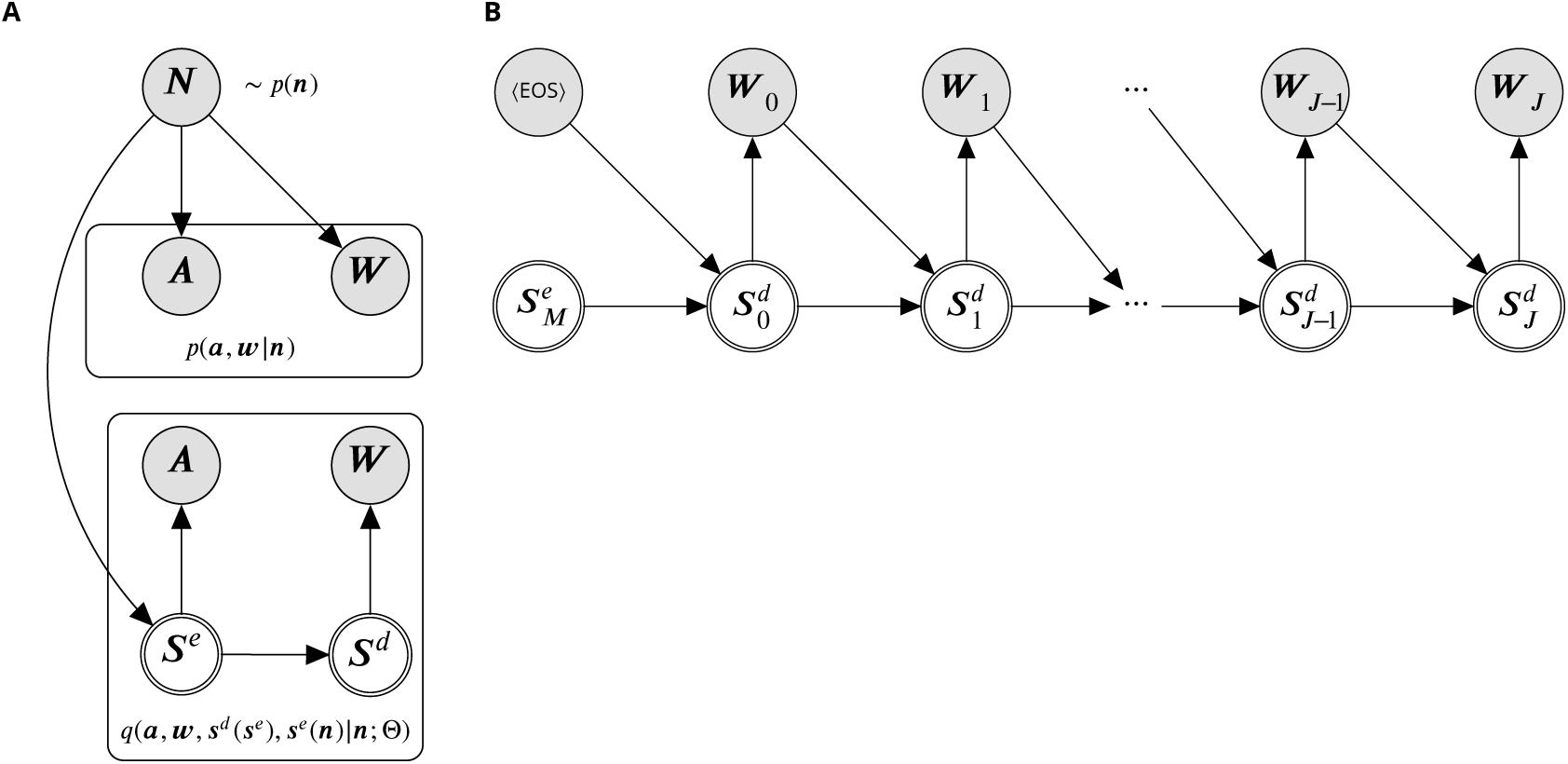
Graphical model for the decoding process. Circles represent random variables; doubled circles are deterministic functions of their inputs. **(A)** The true generative process (above) and the encoder-decoder model (below). The true relationship between neural activity (***N***), the speech-audio signal (***A***), and word sequences (***W***), denoted *p*(***a, w***|***n***), is unknown (although we have drawn the graph to suggest that ***W*** and ***A*** are independent given ***N***). However, we can observe samples from all three variables, which we use to fit the conditional model, *q*(***a, w, s***^*d*^ (***s***^*e*^), ***s***^*e*^(***n***)|***n***; Θ), which is implemented as a neural network. The model separates the encoder states, ***S***^*e*^, which directly generate the audio sequences, from the decoder states, ***S***^*d*^, which generate the word sequences. During training, model parameters Θ are changed so as to make the model distribution *q* over ***A*** and ***W*** look more like the true distribution *p*. **(B)** Detail of the graphical model for the decoder, unrolled in sequence steps. Each decoder state is computed deterministically from its predecessor and the previously generated word or (in the case of the zeroth state) the final *encoder* state and an initialization token, ⟨ EOS⟩.

Similarly, at each step of the *word* sequence, we intepret the (vector) output of the *decoder* RNN, ***ŵ***, as a set of categorical probabilities over the words of the vocabulary:

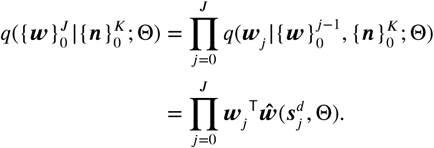

The first equality follows from the chain rule of probability, but the second follows only from the graph in Fig. 5B, and embodies the hypothesis that the decoder state, 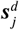, can provide a compact summary of the preceding words in the sequence (i.e., up through step *j* − 1). The second line is consistent with the first because the decoder state 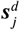 depends on only preceding words and the sequence of neural data, via the recursion

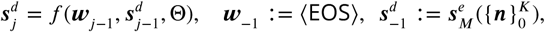

where the function *f* is given by the decoder RNN (see again Fig. 5). Note that the dependence on the neural data enters in only through the final *encoder* state, 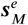. This embodies the hypothesis that all the information about the word sequence that can be extracted from the neural sequence can be summarized in a single, fixed-length vector. In any case, the resulting cross entropy for the word sequences is therefore

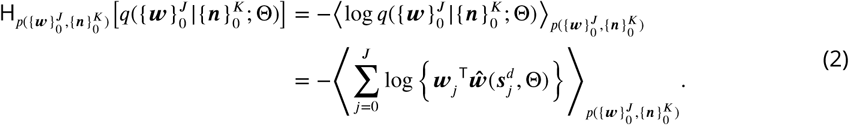

Note that, since the observed words are one-hot, the inner product in the last line simply extracts from the vector of predicted probabilities, ***ŵ***, the predicted probability of the observed word (so that the predicted probabilities of the other words have no effect on the cost function).

The relative importance of the cross entropies in Eqs. 1 and 2 is not obvious *a priori*: ultimately, we require only that the model produce (good) word sequences—no MFCCs need be generated— but MFCC-targeting nevertheless guides the network toward better solutions (especially early in training). In practice, then, we set the loss function equal to a weighted sum of the penalties above (dropping the constants), with the weight, *λ*, determined empirically (see below):

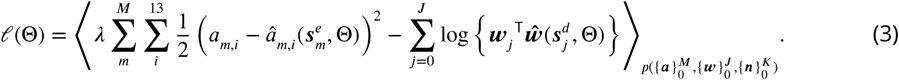

As usual, we minimize this loss by stochastic gradient descent. That is, we evaluate the gradient (with respect to Θ) of the function in brackets not under the total data distribution *p*, but rather under a random subset of these data; take a step in the direction of this gradient; and then repeat the process until approximate convergence.

#### Implementation: the Data

A single training datum consists of the triple 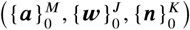. The neural sequence 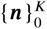 consists of vectors of high-*γ* activity from precisely (see **Data collection and pre-processing**) that period of time during which the participant produced the sequences of words and MFCCs. The length *K* of this sequence thus depends on this time but also on the sampling rate (approximately 200 Hz). Before entering the neural network, this sequence is reversed in time, in order to reduce the number of computations separating the initial element of the input sequence from the (presumably most correlated) initial element of the output (word) sequence (***Sutskever et al., 2014***). The length of each vector in this sequence is equal to the number of (functioning) ECoG channels.

Similarly, the length *J* of the word sequence is simply the number of words in the sentence, plus one extra terminating token, ⟨EOS⟩. A single element of this sequence, ***w***_*j*_, i.e. a “word,” is likewise a vector, being a one-hot encoding, with length equal to the vocabulary size (about 1800 for MOCHA-TIMIT and 125 for the picture descriptions; see Table 3). This includes an out-of-vocabulary token, ⟨OOV⟩, to cover words not in the actual sentence sets but erroneously produced by the participants (in practice less than one percent of the data).

The length *M* of the MFCC sequences would seem, at first blush, to be perforce identical to *K*, the length of the neural sequences, since the encoder neural network maps each element of the input sequence to an output. However, the two layers of temporal convolution that precede the encoder RNN effectively decimate the neural sequences by a factor of twelve (see next section for details). Since the input sequences are initially sampled at about 200 Hz, data thus enters the encoder RNN at about 16 Hz. To achieve the same sampling rate for the audio signal, the MFCC sequences were simply decimated by a factor of twelve, starting from the zeroth sequence element. In fact, the audio sequences ought to be low-pass filtered first (at about 8 Hz) to prevent aliasing, but since the production of high-fidelity MFCCs is not ultimately a desideratum for our network, in practice we used the crude appoximation of simply discarding samples. The length of a single element of the MFCC sequence is 13, corresponding to the total frame energy (first element) and MFCCs 2-13 (see **Speech audio signal** above).

The sequences in any given triple 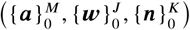 will not in general have the same lengths as the sequences of any other triple, since speaking time and the number of words per sentence vary by example. The network was nevertheless trained in mini-batches, simply by zero-padding the data out to the longest sequence in each mini-batch, and making sure to read out RNN outputs at each sequence’s true, rather than nominal, length (see next section). Clearly, training will be inefficient if mini-batches are dominated by padding, which can happen if (e.g.) one input sequence is much longer than the others. To alleviate this, one can try to group sentences into mini-batches with similar lengths, but we did not attempt such expedients. Instead, we simply enforced a maximum sentence length of 6.25 seconds, which in practice truncated less than one percent of examples. Mini-batches were then created simply by randomizing, at the beginning of each epoch, the order of the sequences, and then dividing the result into consecutive batches of 256 examples.

#### Implementation: Architecture

The basic architecture of the network, shown in Fig. 6, was modeled after the encoder-decoder neural network for machine translation of ***Sutskever et al.*** (***2014***), although there are significant modifications.

**Figure 6.**
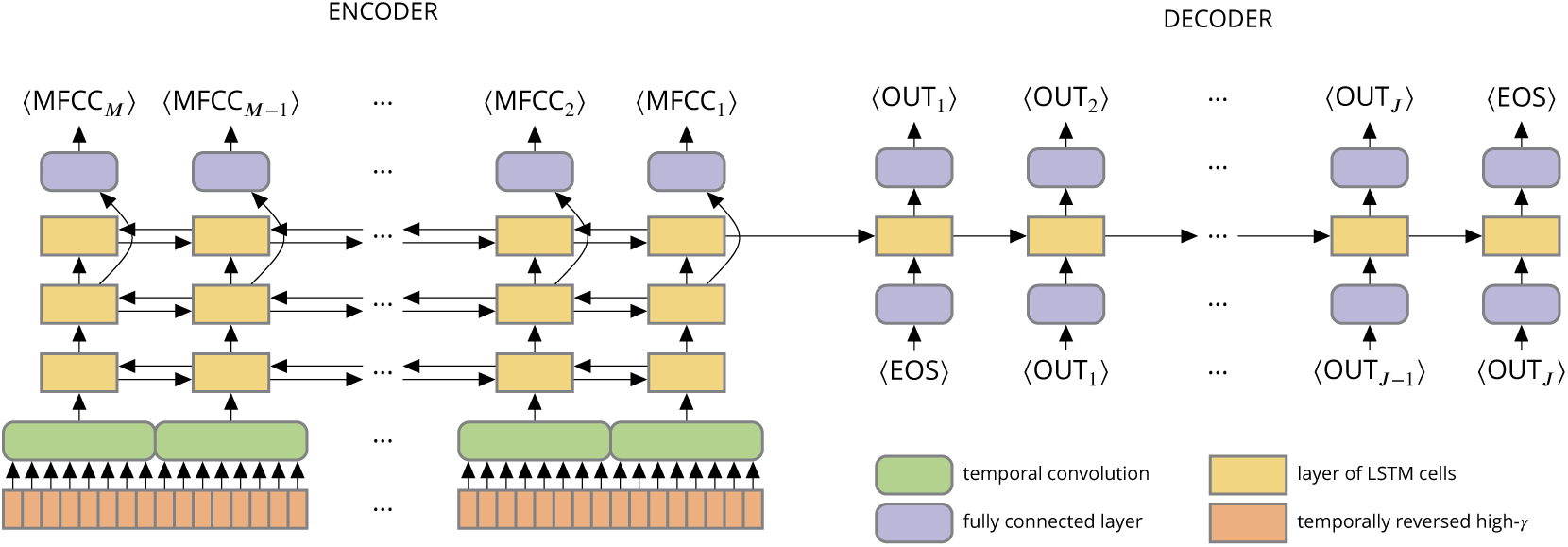
Network architecture. The encoder and decoder are shown unrolled in time—or, more precisely, sequence elements (columns). Thus all layers (boxes) within the same row of the encoder or of the decoder have the same incoming and outgoing weights. The arrows in both directions indicate a bidirectional RNN. Although the figure depicts the temporal convolutions as eight samples wide (due to space constraints), all results are from networks with twelve-sample-wide convolutions. The end-of-sequence token is denoted ⟨EOS⟩.

Each sequence of ECoG data (orange boxes, bottom left) enters the network through a layer of temporal convolution (green boxes). The stride length (i.e., number of samples of temporal shift) of the convolutional filters sets the effective decimation factor—in this case, twelve. In this network the filter width is also fixed to the stride length. This order-of-magnitude downsampling is crucial to good performance: without it, the input sequences are too long even for the LSTM cells to follow. Since the analytic amplitude does not have much content beyond about 20 Hz, the procedure also throws away little information. The convolutional layer consists of 100 filters (“channels”); no max-pooling or other nonlinearity was applied to them.

Output from the convolutional layer at each time step (i.e., a 100-dimensional vector) passes into the *encoder RNN* (gold rectangles), which consists of three layers of bidirectional RNNs. In particular, each “layer” consists of an RNN processing the sequence in the forward direction (receiving input *m* −1 before receiving input *m*) and an RNN processing the sequence in the backward direction (receiving input *m* before receiving input *m* − 1). The outputs of these RNNs are concatenated and passed as input to the next layer. Each “unit” in a single RNN layer is a cell of long short-term memory (LSTM): a complex of simple units that interact multiplicatively, rather than additively, allowing the model to learn to gate the flow of information and therefore preserve (and dump) information across long time scales (***Hochreiter and Schmidhuber, 1997***). We used the LSTM design of ***Gers et al.*** (***2000***). Since both the forward and backward RNNs have 400 units, the total RNN state is an 800-dimensional vector (the state of the LSTM cells in the two directions, concatenated together).

The outputs of the second (middle) layer of the encoder RNN also pass through a fully connected output layer (bluish boxes; 225 linear units followed by rectified-linear functions) and thence through a “fat” (13 × 225) matrix, yielding the MFCC predictions.

The *decoder RNN* (gold rectangles) is initialized with the final state of the final layer of the encoder RNN. (In fact, this state is a concatenation of the final state of the forward encoder RNN with the first state of the backward encoder RNN, although both correspond to step *M* of the input sequence. Thus the dimension of the decoder state is 800 = 400 · 2.) This RNN receives as input the preceding word, encoded one-hot and embedded in a 150-dimensional space with a fully connected layer of rectified-linear units (bluish boxes below). The decoder RNN is necessarily unidirectional, since it cannot be allowed to access future words. The output of the decoder RNN passes through a single matrix (bluish boxes) that projects the state into the space of words, with dimensional equal to the vocabulary size. For the picture-description task, with its small vocabulary, this dimension is 125. For MOCHA-TIMIT, we let the output dimension be 1800 even when training and testing only with MOCHA-1, i.e. the first set of 50 sentences, with its much smaller vocabulary (about 250 words).

This facilitated comparisons with cross-task training, as in Fig. 3 in the main text.

The architecture hyperparameters are summarized in Table 4.

**Table 4.** Architecture hyperparameters. The filter strides in the temporal convolution were set equal to the filter widths, *viz.* 3 and 4 in the first and second layer, resp.

| layer type | # units | connectivity | nonlinearity |
| --- | --- | --- | --- |
| encoder embedding | [(100 × 12)] | temporal conv. | none |
| encoder RNN | [400, 400, 400] × 2 | bidirectional | LSTM (Gers *et al.*, 2000) |
| encoder projection | [225, 13] | full | [ReLU, none] |
| decoder embedding | [150] | full | ReLU |
| decoder RNN | [800] | unidirectional | LSTM (Gers *et al.*, 2000) |
| decoder projection | [1808] or [125] | full | none |

### Training, Testing, Hyperparameter Optimization, and Cross-Validation

#### Training

The network described in the previous section was implemented in TensorFlow, an open-source machine-learning framework with a Python API (***Abadi et al., 2016***). Gradient descent was performed with AdaM optimization (***Kingma and Ba, 2014***). Dropout (***Srivastava et al., 2014***) was applied to all layers, but the network was not regularized in any other way (e.g. weight decay). Dropout in the RNN was applied to the non-recurrent connections only (***Zaremba et al., 2014***).

Across-participant transfer learning proceeded as follows. First, the network was initialized randomly and then “pre-trained” for 200 epochs on one participant. Then the two input convolutional layers were reset to random initial values, all other weights in the network were “frozen,” and the network was trained on the second (target) participant for 60 epochs. That is, the error gradient was backpropagated through the entire network but *only* the convolutional layers were updated. Thus, during this stage of training, the convolutional layers are trained to extract, from the *second* participant’s data, features that work well with an encoder-decoder network fit to the *first* participant’s data. Finally, the weights in the encoder-decoder were unfrozen, and the entire network “post-trained” for another 540 epochs (for a total of 800 epochs). This allowed the rest of the network to accommodate idiosyncracies in the second participant’s data.

#### Testing

To test the network, data are passed through as during training, with one very important, and one minor, difference. During both training and testing, the output of the decoder provides a probability distribution over word sequences:

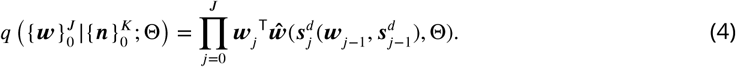

During training, it is necessary to evaluate this distribution only under each *observed* word sequence, 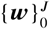. That is, at each step in the sequence, the *output* of the network is evaluated only at the current observed word (***w***_*j*_ in the right-hand side of Eq. 4); and likewise the *input* to the network is set equal to a one-hot vector encoding the previous observed word (***w***_*j*−1_ in the right-hand side of Eq. 4). During testing, however, one would like to compute the probability distribution over *all* sequences, or at least to find the most probable sequence under this distribution. Evaluating all sequences is, however, intractable, because the number of sequences grows exponentially in the sequence length. Instead, we employ the usual heuristic to find the most probable sequence: At each step, we simply pick the most likely word, and use it as input at the next step (***Sutskever et al., 2014***). This is not guaranteed to find the most probable sequence under the model distribution because (e.g.) the first word in this most probable sequence need not be the most probable first word. To alleviate this difficulty, it is possible to maintain a “beam” (as opposed to point) of the *N* most probable sequences, where the width *N* of the beam controls the trade-off between optimality and tractability—but in our experiments using a beam search did not notably improve performance, so we did not use it.

The minor difference between testing and training is in the set of parameters used. We evaluate the network not under the final parameter values, Θ_*T*_, but rather under an exponential moving average of these parameters, 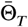, across update steps *t*:

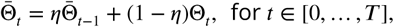

where the decay rate *η* is a hyperparameter. This smooths out shocks in the weight changes.

Training and testing hyperparameters and their values are listed in Table 5.

**Table 5.** Training hyperparameters. RNN = recurrent neural network, MFCC = Mel-frequency cepstral coefficients, EMA = exponential moving average (see text); TL = transfer learning.

| parameter | value |
| --- | --- |
| learning rate | 0.0005 |
| feed-forward dropout | 0.1 |
| RNN dropout | 0.5 |
| MFCC-penalty weight, $\lambda$ | 1.0 |
| examples/mini-batch | 256 |
| EMA decay, $\eta$ | 0.99 |
| # training epochs | 800 |
| TL: # pre-training epochs | 200 |
| TL: # training epochs | 60 |
| TL: # post-training epochs | 540 |

#### Hyperparameter Optimization and Cross-Validation

All hyperparameters were chosen based on performance of a single participant (B, pink in the main-text figures) on a single validation block. Initial choices were made *ad hoc* by trial and error, and then a grid search was performed for the dropout fractions, the MFCC-penalty weight (*λ*), and the layer sizes. This validation block was not, in fact, excluded from the final tests, since it was but one tenth of the total test size and therefore unlikely to “leak” information from the training into the results.

Results (word error rates) were cross validated. For each evaluation, *N* randomly chosen blocks were held out for testing, and a network was trained from scratch on the remaining blocks, where *N* was chosen so as to hold out approximately 10% of the data. Numerical breakdowns are given in Table 3.

#### Significance Testing

Recall that each bar in Fig. 2A and Fig. 3 shows the average (and its standard error) word error rate (WER) across 30 models trained from scratch and tested on randomly held-out validation blocks. This randomization was performed separately for each model, so no two models (bars in the plot) to be compared are guaranteed to have precisely the same set of validation blocks. Nevertheless, the identity of the validation block does influence the results, so a paired test is expected to reduce the variance of differences in WERs. Therefore, in order to employ a paired test, we simply restricted our comparison to the subsets (one from each model) of maximal size with matching validation blocks. To the WER differences from these validation-block-matched pairs, we applied a (one-sided) Wilcoxon signed-rank test, asking whether a particular model or form of transfer learning were not superior to its rivals. The resulting p-values were then Holm-Bonferroni corrected for multiple comparisons. For example, the encoder-decoder network was compared with five other decoders (four variants, plus the phoneme-based HMM), so the p-values reported for Fig. 2A were corrected for five comparisons. The transfer-learning results were corrected for fourteen comparisons: the twelve comparisons annotated in Fig. 3, plus the two comparisons of WER with and without transfer learning for the picture-description data (not shown in the figure but discussed in the main text).

### The Relative Contributions of Electrodes to Decoding

The contribution of an individual electrode, and therefore local anatomical area, might be estimated in multiple ways. Perhaps the most straightforward is simply to train a network with that electrode left out, and measure the increase in word error rate (WER). Unfortunately, that increase will generally be small compared to the variation in WER across retrainings due simply to the randomness of stochastic gradient descent with random initialization, and therefore hard to detect without repeated retrainings—each of which takes upwards of 45 minutes of wall time. Multiplying these 45 minutes by the number of electrodes (about 250) and again by the number of repeats required to detect the WER signal in the noise of retrainings (about 10) yields a prohibitively large amount of computation time.

Alternatively, this electrode-omission procedure could be modified for groups of electrodes, each perhaps corresponing to a gross anatomical area. But even setting aside the loss of spatial resolution, second-order effects—i.e., interactions between (groups of) electrodes—would be ignored. E.g., the electrode-omission procedure would underestimate the contribution of those electrodes that contribute significantly to decoding when present, but for which the network can to some extent compenstate, when absent, by leaning on other channels.

Instead, then, we examine the gradient of the loss function, Eq. 3, with respect to the *inputs*, i.e., the sequences of high-*γ* activity. This measures how much small deviations from an input sequence at each electrode affect the loss, and is the same quantity proposed by (***Simonyan et al., 2013***) to determine which regions of an image are most useful to its classification by a convolutional neural network. In the present case, we should like to know the relative usefulness of electrodes, not for a particular sequence of ECoG data, nor for a particular time in the sequences, but for all sequences at all moments in time. To remove this “nuisance” variation, we take the norm of the derivatives across example sequences and time steps within those sequences. (We use a norm rather than an average because it is the magnitudes of the derivatives that matter: it doesn’t matter whether an increase or a decrease in the high-*γ* activity is required to decrease the loss.) The gradient itself is computed via backpropagation through the trained model, all the way into the testing (as opposed to training) data.

Since we are interested only in relative electrode contributions within, rather than across, subjects, for display in Fig. 4 we rescaled all data into the same range of arbitrary units.

### Phoneme-Based Sentence Classifier

The Viterbi decoders against which the encoder-decoder models were compared (e.g., Fig. 2A) were trained and tested as follows (see also (***Moses et al., in press***)). First, phonetic transcriptions were obtained for each sentence, aligned with the neural data. Next, small time windows of high-*γ* activity around each time point were projected onto their first few principal components, yielding low-dimensional neural features. Finally, a (fully observed) hidden Markov model with Gaussian emissions was trained to map phoneme identities to these neural features. However, rather than learn the hidden-state transition probabilities from the data, and infer phoneme sequences from test data under the resulting model, inferences were made with 50 different transition-probability models, one for each sentence in the MOCHA-1 set. Each model allowed only those transitions consistent with the corresponding sentence (transition to the next phoneme in the sequence, or a self transition). For each of these 50 transition-probability models, the most probable (Viterbi) hidden path and its corresponding probability were computed; and then the sentence corresponding to the most probable path over all models was selected. This process of training and testing was performed ten times with a leave-one-block-out cross-validation scheme to obtain the results shown in Fig. 2A.

## Acknowledgments

The project was funded by a research contract under Facebook’s Sponsored Academic Research Agreement. Data were collected and pre-processed by members of the Chang lab, some (MOCHA-TIMIT) under NIH grant U01 NS098971. Some neural networks were trained using GPUs generously donated by the Nvidia Corporation.

## Supplementary Material

**Figure S1.**
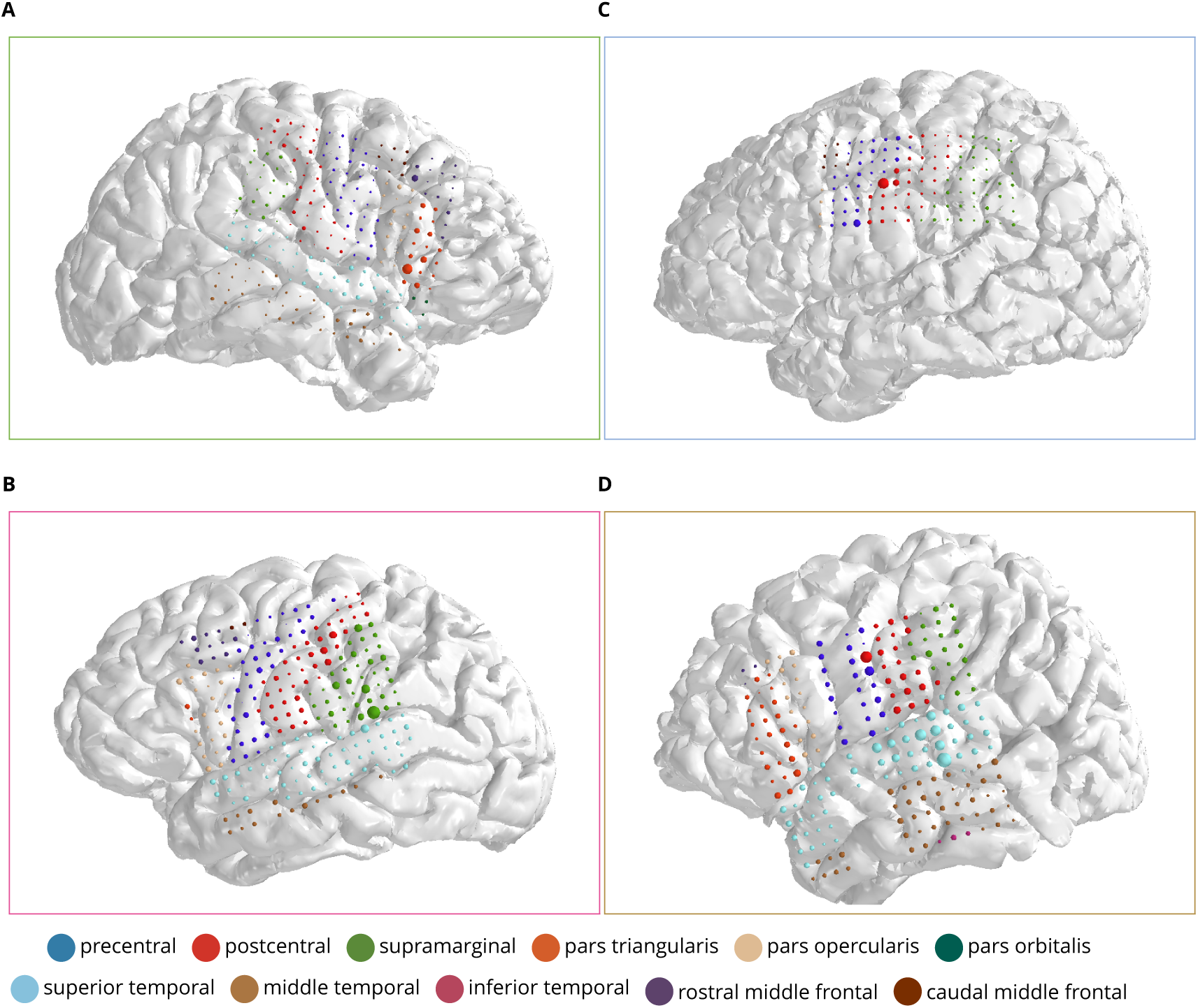
Electrode coverage and contributions. **(A–D)**: Anatomical reconstructions of the four participants (colored frames indicating subject identity according to the color scheme used throughout), with the location of the ECoG electrodes indicated with colored discs. For each disc, the *area* indicates the electrode’s contribution to decoding (see Methods), and the *color* indicates the anatomical region (see legend).

## Footnotes

1 Clearly, the number of words (*J*) in a given sentence will not be the same as the number (*K*) of vectors in a given ECoG sequence, or the number (*M*) in a given MFCC sequence. But neither will *K* and *M* be equal, since MFCCs need to be downsampled relative to the ECoG sequence, due to the decimating effect of the temporal convolution. We discuss this below. Also, to lighten notation, we do not mark the fact that these integers are themselves random variables.

## References

Abadi M, Agarwal A, Barham P, Brevdo E, Chen Z, Citro C, Corrado GS, Davis A, Dean J, Devin M, Ghemawat S, Goodfellow I, Harp A, Irving G, Isard M, Jia Y, Jozefowicz R, Kaiser L, Kudlur M, Levenberg J, et al. TensorFlow: Large-Scale Machine Learning on Heterogeneous Distributed Systems. Google Research; 2016.

Bahdanau D, Cho K, Bengio Y. Neural Machine Translation by Jointly Learning to Align and Translate; 2014, http://arxiv.org/abs/1409.0473v3, doi: 10.1146/annurev.neuro.26.041002.131047.

Bai S, Kolter JZ, Koltun V. An Empirical Evaluation of Generic Convolutional and Recurrent Networks for Sequence Modeling; 2018, https://arxiv.org/pdf/1803.01271.pdf.

Bouchard KE, Mesgarani N, Johnson K, Chang EF. Functional Organization of Human Sensorimotor Cortex for Speech Articulation. Nature. 2013 feb; 495(7441):327–332. http://www.ncbi.nlm.nih.gov/pubmed/23426266, doi: 10.1038/nature11911.

Brumberg JS, Kennedy PR, Guenther FH. Artificial Speech Synthesizer Control by Brain-Computer Interface. In: Interspeech; 2009. p. 636–639. Brumberg2009.

Brumberg JS, Wright EJ, Andreasen DS, Guenther FH, Kennedy PR. Classification of intended phoneme production from chronic intracortical microelectrode recordings in speech-motor cortex. Frontiers in Neuroengineering. 2011; 5(May):1–12. doi: 10.3389/fnins.2011.00065.

Caruana R. Multi-task Learning. Multitask Learning. 1997; 28:41–75.

Cho K, van Merrienboer B, Bahdanau D, Bengio Y. On the Properties of Neural Machine Translation: Encoder–Decoder Approaches. Proceedings of SSST-8, Eighth Workshop on Syntax, Semantics and Structure in Statistical Translation. 2014; p. 103–111. http://arxiv.org/pdf/1409.1259v2.pdf{%}5Cnhttp://arxiv.org/abs/1409.1259

Cho K, van Merrienboer B, Gulcehre C, Bahdanau D, Bougares F, Schwenk H, Bengio Y. Learning Phrase Representations using RNN Encoder-Decoder for Statistical Machine Translation. In: 2014 Conference on Empirical Methods in Natural Language Processing (EMNLP); 2014. p. 1724–1734. doi: 10.3115/v1/D14-1179.

Conant DF, Bouchard KE, Leonard MK, Chang EF. Human Sensorimotor Cortex Control of Directly Measured Vocal Tract Movements during Vowel Production. The Journal of Neuroscience. 2018; 38(12):2955–2966. doi: 10.1523/jneurosci.2382-17.2018.

Dichter BK, Breshears JD, Leonard MK, Chang EF. The Control of Vocal Pitch in Human Laryngeal Motor Cortex. Cell. 2018; 174(1):21–31.e9. https://doi.org/10.1016/j.cell.2018.05.016, doi: 10.1016/j.cell.2018.05.016.

Germann U, Aligned Hansards of the 36th Parliament of Canada; 2001. https://www.isi.edu/natural-language/download/hansard/, accessed: 2018-12-20.

Gers FA, Schmidhuber J, Cummins F. Learning to Forget: Continual Prediction with LSTM. Neural Computation. 2000; 12(10):2451–2471. doi: 10.1162/089976600300015015.

Gilja V, Pandarinath C, Blabe CH, Nuyujukian P, Simeral JD, Sarma AA, Sorice BL, Perge JA, Jarosiewicz B, Hochberg LR, Shenoy KV, Henderson JM. Clinical Translation of a High-Performance Neural Prosthesis. Nature Medicine. 2015; 21(10):1142–1145. doi: 10.1038/nm.3953.

Herff C, Heger D, de Pesters A, Telaar D, Brunner P, Schalk G, Schultz T. Brain-to-Text: Decoding Spoken Phrases from Phone Representations in the Brain. Frontiers in Neuroscience. 2015; 9 (JUN):1–11. doi: 10.3389/fnins.2015.00217.

Hochreiter S, Schmidhuber J. Long Short-Term Memory. Neural Computation. 1997; 9(8):1735–1780.

Jarosiewicz B, Sarma AA, Bacher D, Masse NY, Simeral JD, Sorice B, Oakley EM, Blabe C, Pandarinath C, Gilja V, Cash SS, Eskandar EN, Friehs G, Henderson JM, Shenoy KV, Donoghue JP, Hochberg LR. Virtual Typing by People with Tetraplegia Using a Self-Calibrating Intracortical Brain-Computer Interface. Science Translational Medicine. 2015; 7(313):1–19. http://stm.sciencemag.org/cgi/doi/10.1126/scitranslmed.aac7328, doi: 10.1126/scitranslmed.aac7328.

Kingma DP, Ba J. Adam: A Method for Stochastic Optimization; 2014, http://arxiv.org/abs/1412.6980, doi: 10.1063/1.4902458.

Lyons J, Python Speech Features; 2018. https://github.com/jameslyons/python_speech_features, [Online; version 0.6].

Moses DA, Leonard MK, Makin JG, Chang EF. Real-Time Decoding of Question-and-Answer Speech Dialogue Using Human Cortical Activity. Nature Communications. in press;.

Mugler EM, Tate MC, Livescu K, Templer JW, Goldrick MA, Slutzky MW. Differential Representation of Articulatory Gestures and Phonemes in Precentral and Inferior Frontal Gyri. The Journal of Neuroscience. 2018; 4653:1206–18. http://www.jneurosci.org/lookup/doi/10.1523/JNEUROSCI.1206-18.2018, doi: 10.1523/JNEUROSCI.120618.2018.

Munteanu C, Penn G, Baecker R, Toms E, James D. Measuring the Acceptable Word Error Rate of Machine-Generated Webcast Transcripts. In: Interspeech; 2006. p. 157–160.

Nuyujukian P, Albites Sanabria J, Saab J, Pandarinath C, Jarosiewicz B, Blabe CH, Franco B, Mernoff ST, Eskandar EN, Simeral JD, Hochberg LR, Shenoy KV, Henderson JM. Cortical Control of a Tablet Computer by People with Paralysis. PLoS ONE. 2018; 13(11):1–16. doi: 10.1371/journal.pone.0204566.

Pei X, Barbour DL, Leuthardt EC. Decoding Vowels and Consonants in Spoken and Imagined Words Using Electrocorticographic Signals in Humans. Journal of Neural Engineering. 2011; 8(4):1–11. doi: 10.1088/1741-2560/8/4/046028.

Pratt L, Mostow J, Kamm C. Direct Transfer of Learned Information Among Neural Networks. AAAI-91 Proceedings. 1991; p. 584–589. www.aaai.org.

Rumelhart D, Hinton GE, Williams RJ. Learning Representations by Back-Propagating Errors. Nature. 1986; 323(9):533–536. doi: 10.1139/p72-256.

Schalkwyk J, Beeferman D, Beaufays F, Byrne B, Chelba C, Cohen M, Garret M, Strope B. Google Search by Voice: A Case Study. In: Neustein A, editor. Advances in Speech Recognition rpinger; 2010.p. 61–90.

Simonyan K, Vedaldi A, Zisserman A. Deep Inside Convolutional Networks: Visualising Image Classification Models and Saliency Maps; 2013, http://arxiv.org/abs/1312.6034.

Srivastava N, Hinton GE, Krizhevsky A, Sutskever I, Salakhutdinov RR. Dropout: A Simple Way to Prevent Neural Networks from Overfitting. Dropout: A Simple Way to Prevent Neural Networks from Overfitting. 2014; 15:1929–1958.

Stavisky SD, Rezaii P, Willett FR, Hochberg LR, Shenoy KV, Henderson JM. Decoding Speech from Intracortical Multielectrode Arrays in Dorsal “Arm/Hand Areas” of Human Motor Cortex. Proceedings of the Annual International Conference of the IEEE Engineering in Medicine and Biology Society, EMBS. 2018; p. 93–97. doi: 10.1109/EMBC.2018.8512199.

Sutskever I, Vinyals O, Le QV. Sequence to Sequence Learning with Neural Networks. In: Advances in Neural Information Processing Systems 27: Proceedings of the 2014 Conference; 2014. p. 1–9. doi: 10.1007/s10107-014-0839-0.

Szegedy C, Liu W, Jia Y, Sermanet P, Reed S, Anguelov D, Erhan D, Vanhoucke V, Rabinovich A. Going Deeper with Convolutions. In: 2015 IEEE Conference on Computer Vision and Pattern Recognition (CVPR) IEEE; 2015. p. 1–9. doi: 10.1109/CVPR.2015.7298594.

Tian X, Poeppel D. Mental Imagery of Speech and Movement Implicates the Dynamics of Internal Forward Models. Frontiers in Psychology. 2010; 1(OCT):1–23. doi: 10.3389/fpsyg.2010.00166.

Wrench A, MOCHA-TIMIT; 2019. http://www.cstr.ed.ac.uk/research/projects/artic/mocha.html, online database.

Xiong W, Droppo J, Member S, Huang X, Seide F, Seltzer ML, Member S, Stolcke A. Toward Human Parity in Conversational Speech Recognition. IEEE/ACM Transactions on Audio, Speech, and Language Processing. 2017; 25(12):2410–2423.

Zaremba W, Sutskever I, Vinyals O. Recurrent Neural Network Regularization; 2014, http://arxiv.org/abs/1409.2329{%}5Cn http://www.arxiv.org/pdf/1409.2329.pdf.

